# Study of aging using circuit tests and parametric analysis

**DOI:** 10.1101/2022.03.24.485723

**Authors:** Tomokazu Konishi, Shuichi Yanai, Shogo Endo

**Affiliations:** Akita Prefectural University; Tokyo Metropolitan Institute of Gerontology

## Abstract

Mouse development and aging were investigated in 11 tests. The test data were parametrically analysed, normalized, and integrated using principal component analysis (PCA). Changes in the indexing items displayed two trends: power and accuracy in the trials. Power decreased monotonically with age, whereas accuracy peaked at 6 months. Features, such as fear or anxiety, could not have been measured. The noise level and sensitivity of each test were determined. In an open field test that involved a video recording of movement, the noise level was the lowest and the sensitivity was high. This test contributed the most to the PCA axes, and reproduced the two main directions.

**Significance Statement:** The mathematical aspect may have been neglected in behavioural tests. Here, the circuit test data were objectively analysed by focusing on the data distributions during the aging of mice. The test results revealed two clear directions for the data: power, which was gradually reduced, and accuracy, which peaked at sixth months old. Compared with the open field test evaluated using video recordings, the other tests had higher noise levels and weaker signals. Other tests can be improved by using movie recordings and identifying appropriate items.

## Introduction

Various practices have been devised to study the behaviour of laboratory animals. Each method is considered to measure a characteristic property and the results represent different indexing items (Whishaw and Kolb, 2004; Quillfeldt, 2016). Analytical methods of data have been developed to enable efficient testing; for example, many factors are considered in experiments such as the open-field tests (Walsh and Cummins, 1976; Goma and Tobena, 1978; Kafkafi and Elmer, 2005; Stanford, 2007; Ennaceur, 2014; Tatem et al., 2014; Quillfeldt, 2016; Belovicova et al., 2017). Test animals are employed to identify items that serve as indexes for specific behaviours. By using C57BL/6J mice, Yanai and Endo (2021) evaluated the changes in the aging index using different measures. Based on their results, aging is not a linear process, but is instead an asynchronous complex process. Of note, this property has been repeatedly revealed (Lione et al., 1999; Kirkwood, 2005; de Magalhães et al., 2012; Rando and Wyss-Coray, 2021).

On the other hand, whether these measurements represent the factors that need to be studied remains an unanswered question (Walsh and Cummins, 1976; Stanford, 2007; Ennaceur, 2014). For example, the marble burying test is widely used as an index for anxiety; however, the interpretation and validity of its results are controversial (Borsini et al., 2002; Wolmarans et al., 2016); this is partly because the data obtained via these measurements and their relation to reality are difficult to interpret.

Here we wish to point out a problem in data analysis. To date, the accuracy of the analytical methods used in the present study has been questionable as the resulting data are analysed without considering their characteristics. An example of this can be viewed in a non-parametric manner, in which data are often analysed. Although a non-parametric model can be flexibly applied to a wide range of data, such complex model has a falsifiability problem: are the assumptions that form the basis of the model appropriate (Ellis and Silk, 2014; Thornton, 2018). If these assumptions are inappropriate, the accuracy of the analysis decreases. In addition, weak falsifiability reduces the objectivity of the analysis.

A parametric approach is more appropriate as it confirms the data distribution of all test results. This approach investigates the characters of data following the completion of all experiments. This approach is called exploratory data analysis (EDA) (Tukey, 1977). The purpose of EDA is to clarify the structure hidden in the data and maximize the information that can be obtained from the data. In fact, when EDA was employed to the video recordings of the open field test, the data showed fairly complex mathematical properties (Konishi and Ohrui, 2020). By recognizing such characteristics, the accuracy of data analysis may be critically improved. In addition, each test is performed to achieve a goal, and the measurements are assigned empirical meanings; however, the validity of those meanings has only been understood to some extent. These meanings would also be confirmed.

In this study, we reanalysed the results of Yanai and Endo (2021) and achieved greater precision; we opted to perform EDA of the circuit test results. Further, the results of the 11 types of trials were integrated to verify whether each trial type could measure its unique characteristics.

The distribution pattern of the data was verified using quantile-quantile (QQ) plots. This method compares the data and standards of a particular distribution using the same quantiles and can rigorously identify each distributional style compared with more commonly used histograms. Knowing the distribution style enables the identification of an appropriate model for data handling and direct comparison of the data with different configurations, as discussed below. In addition, the noise level of each test was measured and compared. In such tests, the noise would be introduced by individual differences and measurement errors; these noises appear in the differences in measurements of an item at each time point. If the noise is normally distributed, its magnitude can be estimated by the standard deviation (SD) of the data. Naturally, this normality of noise is confirmed using the QQ plot. To compare the magnitude of the noise component, regardless of the magnitude of the measured value, the coefficient of variation (CV) = SD/mean is used. The CVs differed significantly, which indicates significant differences in the accuracy of the measurements and implies that each test data point should be appropriately weighted based on the noise level.

The trial results were converted into a mutually comparable state. All data were transformed to achieve a normal or near-normal distribution. Further, the location *μ* and scale *σ* parameters for the data were set as 0 and 1, respectively, by z-normalization. Using *σ* estimated from each month, the normalized data had the same level of noise among the tests. The normalized data were then averaged over repeated measurements of multiple animals to obtain a representative value for each time point in each trial. Finally, a matrix dataset was constructed with time points by items; this can be regarded as a typical multivariate data.

The normalized matrix data consists of many items, which are independent of each other. In a mathematical sense, they have the same number of multiple dimensions as the item and are thus, difficult to comprehend. Accordingly, the matrix was summarized using principal component analysis (PCA). PCA is a method of grouping the same direction of change for multiple items, with a score obtained for each direction; it is frequently used to reduce the multiple dimensions of matrix data (Jolliffe 2002). If multiple tests change in the same manner, the results will be combined, and the complex aging process (Yanai and Endo, 2021) will be organized within this framework. If any of these processes share a common time-course pattern of increase or decrease, PCA will display the data using an axis for each pattern. As a result, the items can be grouped together and the results of these circuit tests can be integrated.

## Materials and Methods

### Tests and animal feeding

Animals were maintained and behavioural tests were performed as described by Yanai and Endo (2021). Although several tests were performed using existing methods, data collection for the open field test was performed using the parametric method and video recordings (Konishi and Ohrui, 2020).

### Experimental Design and Statistical Analysis

Twelve tests were independently performed using 22 to 24 mice (ages, 3, 6, 12, 18, and 22 months). The training effects were also measured in some tests. For each of the five series of monthly records, data distribution was verified using QQ plots, which compare quantiles of data with one of the distribution models on a scatter plot. If the model fits the data distribution, the plot shows a linear relationship. Of note, the data were normalized, as described in the next section. Further, differences between months were compared using box plots and tested using Tukey’s HSD (Tukey, 1949).

### Coefficient of variation (CV)

At each time point of the test, the standard deviation (SD) was robustly estimated using the median absolute deviation (MAD). These values were averaged and employed as the square root of the mean of the square MADs. The mean of the test was robustly estimated using the trimmed mean (trim =0,3). Further, CV was determined as follows: CV = (averaged MAD) / (trimmed mean). Thereafter, the CVs were sorted according to their medians. Such robust estimators will help to escape the effects of outlying observations, which may be caused by some accidents on the animals.

### Normalization of data

Many data points are distributed either normally or log-normally. Log-normal data points were transformed to normal data points using log(*x*_i_ − *γ*), where *x*_i_ is the data and *γ* is the background parameter, ultimately generating a linear relationship with normal distribution. Some data points obeyed the exponent of the exponential distribution (Konishi and Ohrui, 2020), indicating that the logarithms of the data are linear to the logarithms of the exponential distribution. These data were approximated to normal distribution by deriving the square roots. The converted data were then used for box plots.

The transformed data were further normalized to N(0,1) normal distribution by setting the z-score using (*x*_i_ - *μ*) / *σ*; this is the most basic method known as z-normalization. The z-scores are widely used to compare data in different or without units of measure.

It should be noted that the SD of all data was not used as *σ*; this is because the data increased and decreased from month to month. As a result, when the data are combined at one time, they comprise five peaks, which leads to an unwarranted large σ value. Hence, the scale was estimated as the SD obtained from each time point of an item; incidentally, this represents the noise level of the item conveniently. The SDs were averaged for all months using the same value employed to estimate the CVs with MAD. The location *μ* was found to be the trimmed mean (trim = 0.3) of all time points. Here, in particular, z-scores indicate signal/noise, unifying the noise level among items. Hence the values are comparable among tests. However, the range of z-scores may differ among the tests; higher sensitivity and lower noise will expand the range and vice versa. From repeated measurements of multiple animals, a representative value was determined for each time point of an item; this value was the trimmed mean of the z-scores.

The water maze and rotarod tests are repeated trials for one test; the scores improved following training and reached a certain plateau, which was used for the comparisons. The learning effect was estimated as the difference between the start and plateau of each animal.

### PCA

The total number of representative values of the items measured and normalized in each of the 11 tests was 60. These representative values created a matrix of 5-time points × 60 items. This matrix was applied to the singular value decomposition of PCA (Jolliffe 2002). As the data were normalized, they were centered and scaled. There are two related PCs: PC_sample and PC_item (Konishi, 2015). PC_sample presents the score of the representative values of samples at a given time point while PC_item indicates how an item contributes to PC_sample. For example, when PC_sample is monotonically decreasing in a time series, items that decrease have a positive PC_item value, and vice versa. These PCs were scaled for different numbers of items and samples (sPC) and used for the presentation.

## Results

### Distribution models

Many of the identified items followed a normal distribution. Items, such as the velocity of swimming and counts of certain behaviours, as well as the weight and food uptake, obeyed this model (Figs. 1A, S1, S3, S4, S6, S7, S8, S9, and S12).

**Fig. 1.**
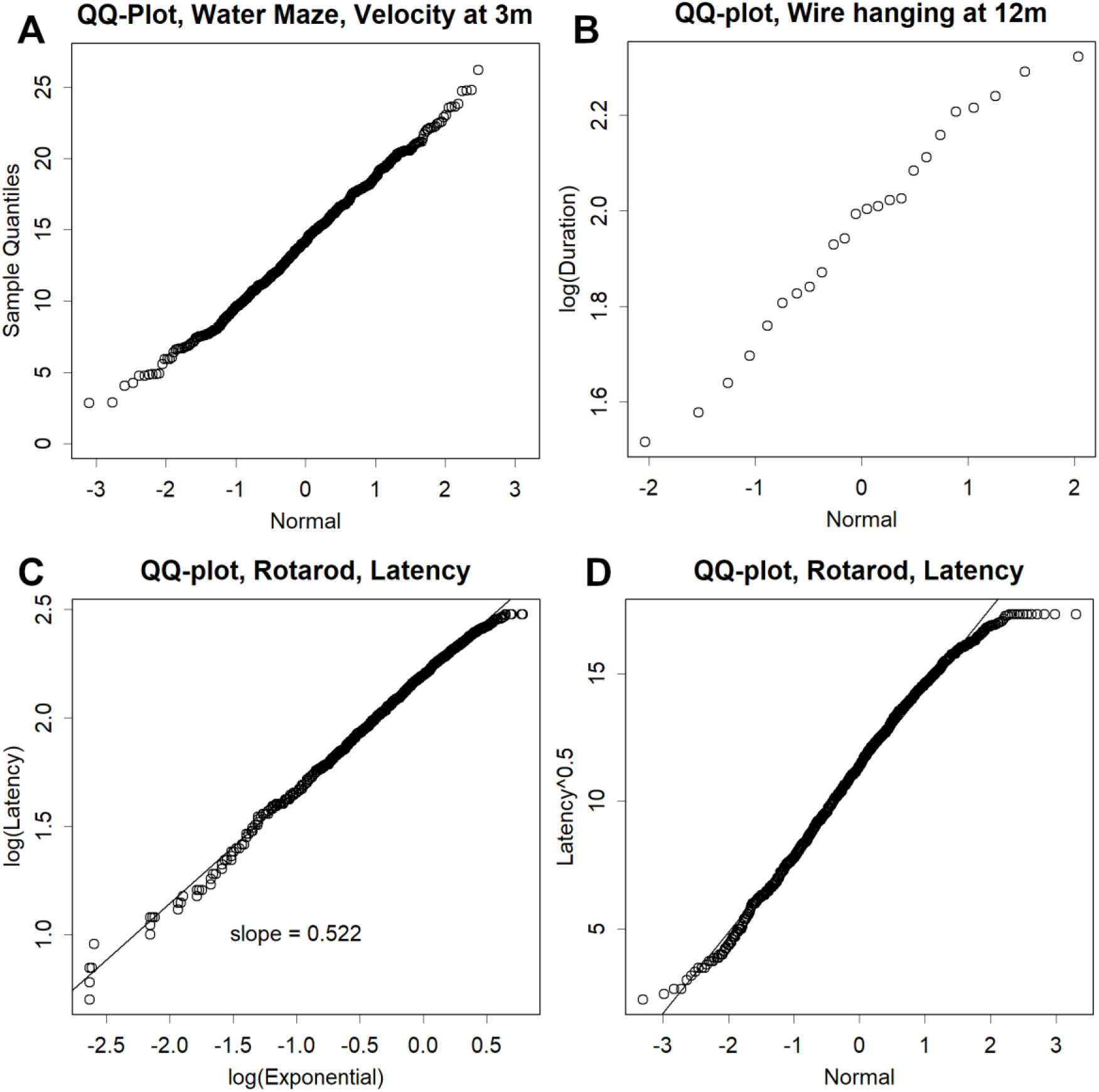
Data distribution in the QQ plots. Typical examples of QQ plots. **A**. Normal distribution of the velocities measured in the water maze test at 3 m. The quantiles of the data were compared with those of the normal distribution. **B**. Lognormal distribution. The logarithms of the duration times measured in the wire hanging test. As the number of data points was low, the plot became discrete. **C**. Exponence of the exponential distribution found in the latency of the rotarod test. Logarithms of the latency quantiles were compared with the logarithms of the exponential distribution. **D**. Example of normalization. The square root of the same data in C was compared with the normal distribution. This simple transformation transferred the data well to near normal conditions.

Other items, such as the learning effect on spatial memory and wire hanging, obeyed the lognormal distribution (Figs. 1B, S2, S3, S7, S8, S9, S12, and S13). Some items may include margins. For example, termination of the water maze and Barnes maze requires a certain period for migration from the start to the endpoint (Figs. S7 and S8). This period is a margin; even the fastest animal cannot have a zero value. As the smallest (and most crowded) section of the lognormal distribution is zero, such margins must be subtracted; otherwise, the distribution will be skewed. Margins were estimated using the background parameter *γ* of the three-parameter lognormal distribution model.

The exponent of the exponential distribution, which has been reported in the open-field test (Konishi and Ohrui, 2020), was also found in rotarod and fear conditioning (Figs. 1C, S3, and S9). This model determines a linear relationship between the logarithms of the data and the logarithms of the exponential distribution. The latencies of certain behaviours and velocities related to latency obeyed this model. These data were normalized by deriving the square roots; this approximation mimicked the normal distribution well (Fig. 1D).

### Training effect

By repeating the trial, the rotarod scores improved and reached a plateau, indicating better motor conditions (Figs. 2A and S5). In addition, scores of latency and distance in the water maze and Barnes maze were improved, highlighting the effect of learning (Figs. 2B, 2C, S7, and S8). In contrast, running (Fig. 2D and S7) and swimming (Fig. S8) were not improved by training. Mice may walk daily in their cages to maintain muscle strength. However, the battery group swam faster than the water-maze only group (Fig. S8), suggesting that variations in training may be helpful for the maintenance of physical strength. The effect of training decreased with age; however, some improvements were observed in month 22 also.

**Fig. 2.**
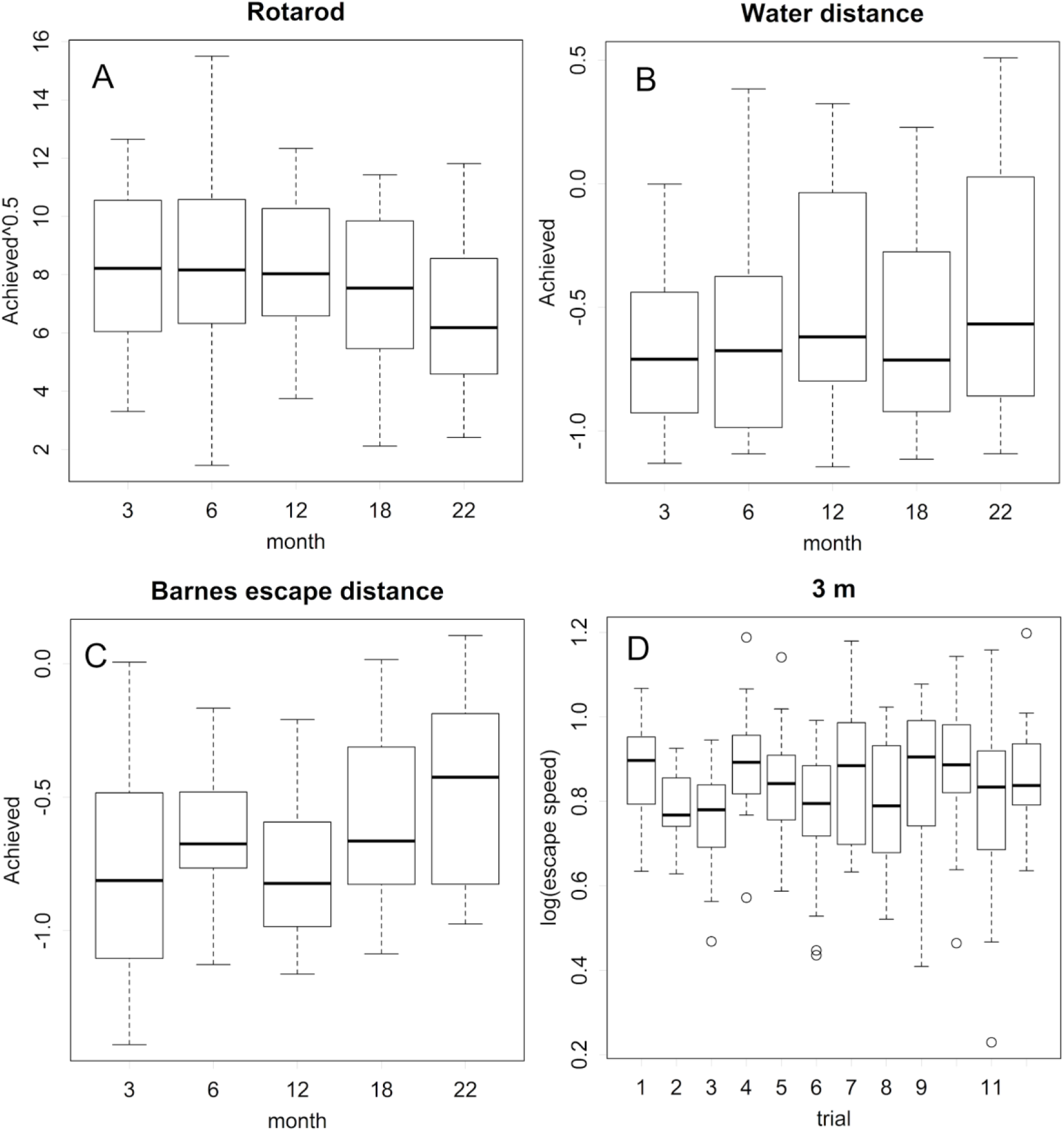
Learning effect by training. **A**. Achievement by training in the Rotarod test. The differences between the plateau and the start of training for each individual are presented. The increase becomes small at 22 months old. **B, C**: Distance travelled to the reward in the water maze and Barnes maze tests. The rate of time savings by the training decreases with age. **D**. Changes in speed in the Barnes maze test at 3 months old. A remarkable change is not obtained by repeating the tests. For other data, see Supplements.

### Noise

The CV of each item reflects the sum of measurement errors and individual differences, and is normalized to derive the differences in magnitude of the measured values, enabling comparable results among tests. The measured items showed exponential differences (Fig. 3) and were distributed lognormally (Fig. S13). The z-scores of log(CV) are presented for easier comparison in each of the supplemental figures.

**Fig. 3.**
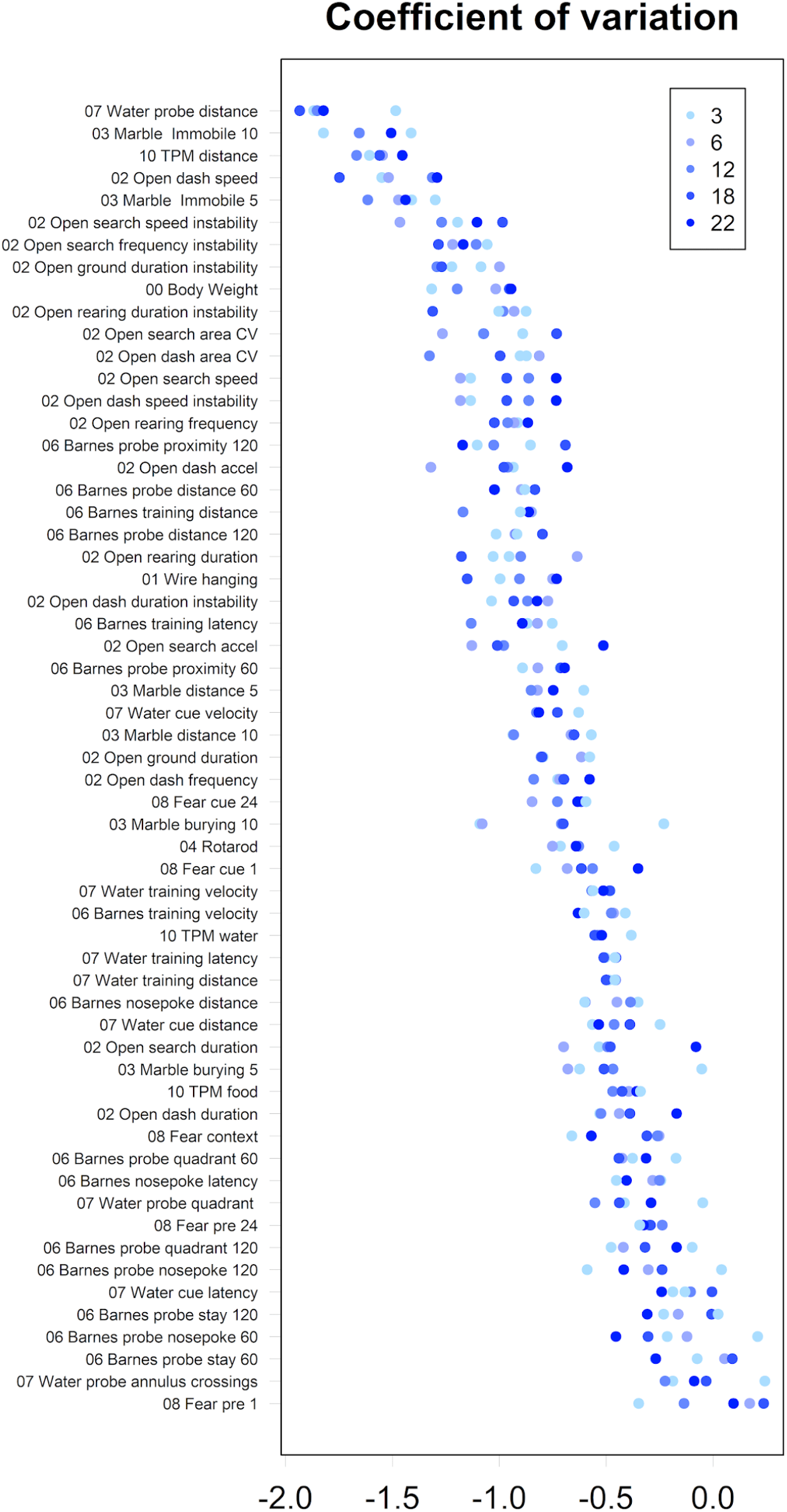
Log of CV for each measurement item. The CV represents the noise level of the individual measurements, which differed on a logarithmic scale. Most of the measurement items had a higher noise level than the weight, which was ninth from the top. Measurements with particularly low CV values were obtained in the open field.

In some tests, only the time to complete a task or count certain behaviours was measured. In these tests, the amount of data to be analysed becomes small, ultimately compromising the reliability and accuracy of the measurements. Additionally, some items may be ineligible from a mathematical perspective. For example, the count of the number of longer stills has been used in open-field tests; however, this is not mathematically appropriate (Konishi and Ohrui, 2020).

Distance tended to have a lower CV than the other items. For example, the CV for distance was markedly lower than that of annulus crossing in the water maze. Many items with lower CV were found in the open field test. Although training did not change the CV, it was found to improve the scores.

The CV increased with age (Fig. 3, deep blue, Fig. S13: box plot). As the test methods were constant, measurement errors were not altered. Such increase may indicate an expansion of individual differences.

### Changes found in development and aging

The changes were summarized by PCA, with two major directions identified: 80% of the data differences appeared at PC1 and PC2 (Fig. S14, Contribution), which would represent the mainstream of development and aging. Fig. 4 shows PC_sample and PC_item on the same graph scale. The PC_sample is a time series, whereas the x-axis of the PC_item has no meaning but is presented in three columns for readability.

**Fig. 4.**
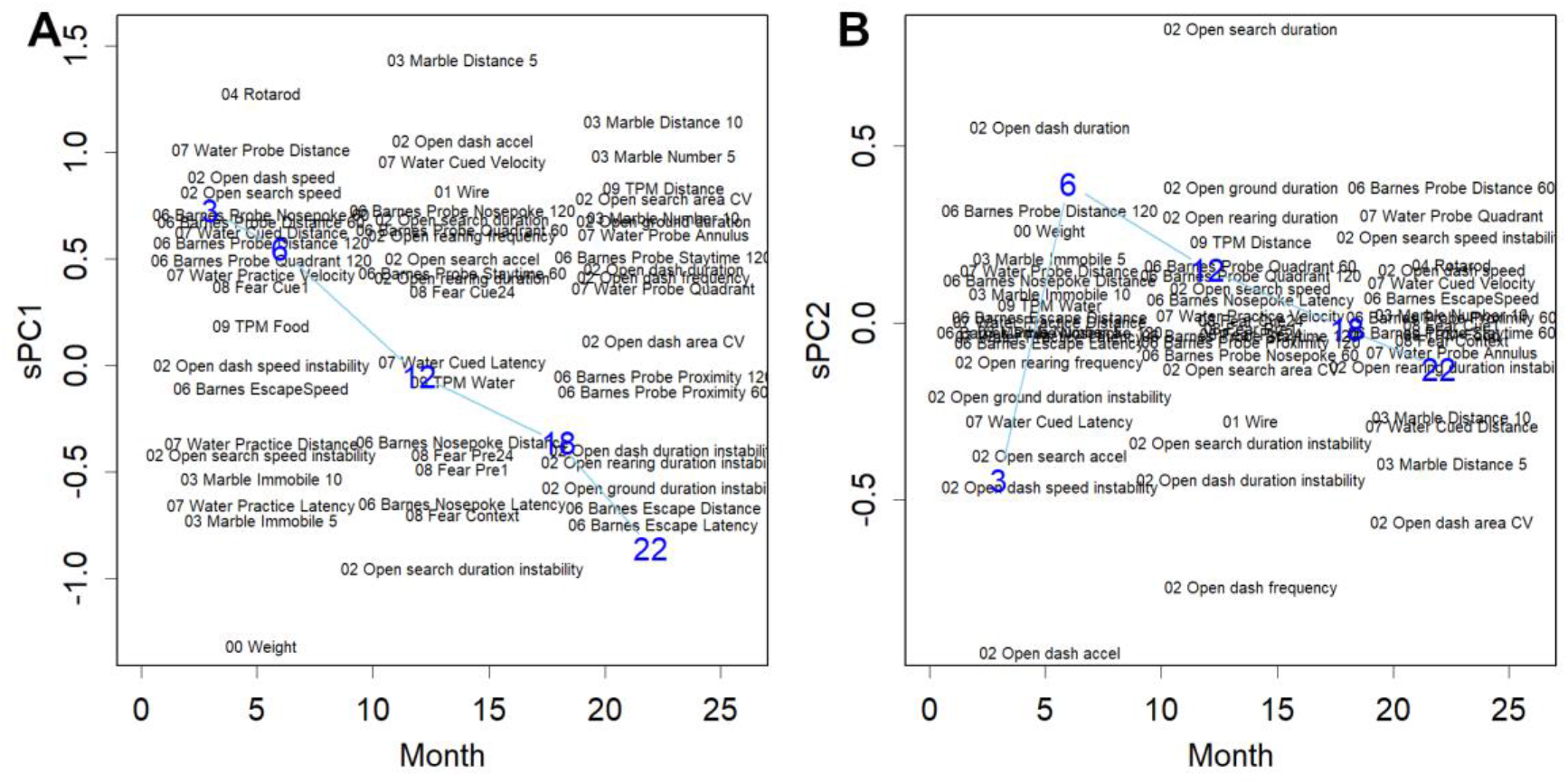
Result of PCA (biplot). Blue numbers indicate the PC for the samples of each age group. **A**. PC1: PC_sample decreased with age (blue). Black letters represent PC_items on the y-axis. The positions on the x-axis for the items are not meaningful, but are displayed in three columns for readability. Items shown above or below indicate those with higher sensitivity. For example, the traveling distance observed in the marble test continued to decrease in the same direction as PC_sample, and was thus shown at the highest position. In contrast, body weight is shown at the lowest position as it continues to increase. **B**. PC2: In the PC_sample, a peak occurs at 6 months old (blue). Refer to the supplement for detailed values, and PC3 and 4.

PC1 included changes that monotonously decreased or increased from month 3 to 22 (Figs. 4A and S14, Table S2: PC_items). Items, such as speed, acceleration, and distance, appeared in this PC, indicating muscle strength as a factor. More power results in a higher acceleration, speed, and distance, which is beneficial for marble hiding and may be related to the increase in weight; older animals do not possess enough power to accelerate their body weight. In addition, items related to learning effects were included, such as those found in the Barnes maze and the water maze. Shorter responses to cues under fear conditions may be included as an effect of learning.

PC2 increased and decreased with a peak at month 6 (Fig. 4B). The items found in the open field had the highest scores. In the peak, animals displayed longer durations of behaviors, such as dash and search. The instability of the speed and acceleration decreased, indicating that they repeated the steady pattern for a longer time; this could be the reason for the long distances in the Barnes maze. However, the frequency of dashes decreased, which suggests better fitness to the purpose of activities with prolonged concentrations.

PC3 and PC4 were concentrated in a small number of items that showed peaks at months 12 and 18, respectively. Both contributed approximately 10% of the data differences; hence, the biplot became narrower (Fig. S14). The major values were recorded for some subjects in the Barnes maze and open field test (Table S2: PC_item).

### The superiority of the open field test

A desirable test should have a lower noise and higher sensitivity for aging. These characteristics can be estimated from the CV and magnitude of PC_item. The open-field test showed superior scores for both noise and sensitivity (Figs. 3 and 4). In particular, this test had higher sensitivity in PC2 than the other tests. This test even could reproduce the shape of PCA alone (Fig. S14).

## Discussion

The data distribution mainly consisted of three patterns, which may be due to the background of the data. Normal distribution (Fig. 1A) represents the sum of random numbers, and appears when the items are determined as the sum of several factors. Many other distributions can be approximated using this model with acceptable accuracy. In addition, when multiple factors produce synergy, they are lognormally distributed, as found in the transcriptome (Konishi, 2005) (Fig. 1B). However, “exponent of the exponential distribution” frequently appears where duration is concerned (Fig. 1C), but is difficult to explain. Originally, the intervals of the randomly occurring phenomena followed an exponential distribution. When a purpose is determined by the synergy of multiple factors, this distribution may be formed. However, the events may not actually occur in a random manner. In fact, in some cases, they occur for a certain purpose, and in other cases, the frequency is altered through the test. The slope of the linear relationship with the distribution model, *a*, indicates the randomness or instability of the behavior; the occurrence is random when *a* = 1, assumes a steady pattern when a < 1 (Fig. 1C), and can be separated into some patterns when *a* > 1 (Fig. S9).

In some tests, the data were far from normal (Fig. 1B and 1C). As non-parametric models often estimate the normality of data, applying such models can artificially expand the noise level of the tests. Inappropriate analyses may have been one of the reasons for the low reliability of these tests (Walsh and Cummins, 1976; Stanford, 2007; Ennaceur, 2014). The accuracy of the analysis can be improved by using proper models; this can be achieved by normalizing the data of a parametric test, such as ANOVA, beforehand.

In many of the tests, the main cause of the noise may not be individual differences, but measurement errors. Although distance cannot be a robust indicator from a mathematical viewpoint (Konishi and Ohrui, 2020), this item had a lower CV than the other items (Fig. 3), as measured in the water maze. Such finding simply suggests the shortcomings of the measuring methods for other items. Of note, the CV values differed on a logarithmic scale. If an item has *n* times the CV, *n*^2^ times of animals will be required to maintain the accuracy of the measurement; however, this is not a practical solution. The measurement method should be improved to reduce noise. If a video recording of the water maze was obtained to evaluate other items, such as speed, better results than those obtained for distance may have been achieved.

The two types of methods used for recording the data showed a higher noise level (Fig. 3). Method one involves counting rare events, such as foot shock, nose poke, and annulus crossing; this inevitably produces discrete numbers with lower resolution. Therefore, the precision of the measurements was limited. Marble burying yielded a better score as the number of marbles prepared for the test was sufficiently large. The second inferior type measures a single duration or arrival time. For example, the duration until falling off was recorded in the rotarod test. If a mouse accidentally fell, this fall would directly influence the results; the same is true for the latency of behavior, such as in the water maze.

The reason for the superiority of the open field test is obvious in the principle of measurement, which is based on movie recordings (Fig. 3 and 4). As each item is determined from a large amount of data, outliers can be removed by robust calculations to avoid the effects of errors. Accordingly, the noise becomes markedly lower than that of the discrete or single measurements of other tests. The accuracy of other tests will improve if movies are recorded to enable in-depth analysis of each movement and identification of items in a statistically valid manner.

Rodent behaviour has been studied in detail, and each test is designed to measure specific behaviour (Quillfeldt, 2016; Belovicova et al., 2017). However, the validity of the tests may not have been examined mathematically. In fact, the comprehensive textbook consistently lacks these statements that relate to mathematical aspects of testing (Whishaw and Kolb, 2004). Noise may have been ignored; however, experiments cannot be objective unless the measurement is shown with the noise level. Evidently, some noisy tests should reassess the measurement method or test itself.

The power and learning effects appeared on the same PC, PC1, as they decreased in the same manner (Fig. 4A); this could occur with or without a correlation between these two factors. It is possible that they were altered in the same manner as the test mice appeared normal without any specific brain problems. In addition, with more time points measured, the improved resolution could separate these changes.

The longer duration of behaviour found in PC2 does not necessarily indicate better endurance. In fact, scores in rotarod or wire hanging, both of which may reflect endurance, did not improve at month 6. Instead, these items may depend on brain function that is required to identify the most suitable actions; this function was not fully developed at month 3. After the peak for month 6, this ability gradually diminished (Fig. 4B).

Aging are being measured via several different methods that are designed to measure different characteristics. Following observation, aging was not found to be linear but instead followed multiple paths (Yanai and Endo, 2021). However, when the results were normalized and summarized by PCA, most of the data converged in only two directions, with PC1 and PC2 representing power and accuracy, respectively. The remaining PC3 and PC4 summarized data from months 12 and 18 as peaks; however, only few items were related and many were found in the open field test. Therefore, regardless of the purpose of the test design, many tests failed to reveal the unique features of aging. This finding may be due to several reasons. First, if we measure more dates, we may be able to obtain more resolution and identify more directions of change. In addition, if a more specific situation is studied rather than simple aging, other directions could appear. However, many tests may inappropriately collect data and noise level is too high to achieve their intended purpose. Here we found no evidence that emotional processes, such as anxiety or fear (Belovicova et al., 2017), have been measured; rather, the methods served as another approach to measure power and accuracy.

Extended Database is available in Figshare at https://doi.org/10.6084/m9.figshare.19417292.v1

